## Supplemental figures S1_S12 for "Identification of a stereotypic molecular arrangement of endogenous glycine receptors at spinal cord synapses"

### Supplementary Information

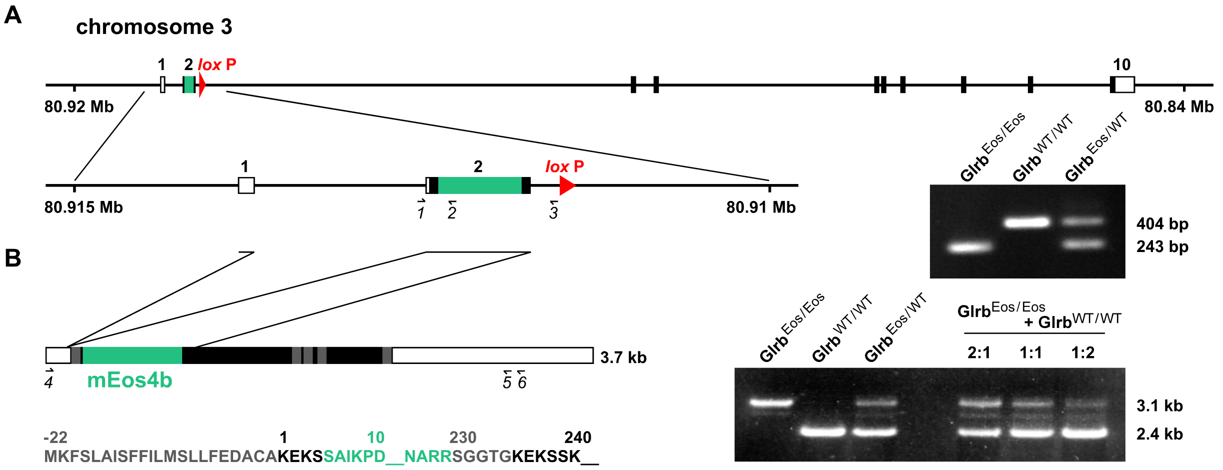


**Fig. S1. Generation of the mEos4b-GlyRβ knock-in mouse model.**

(A) The coding sequence of mEos4b (green) was inserted in exon 2 of the *Glrb* gene by homologous recombination, as shown by the amplification of a 243 bp PCR product in genomic DNA from *Glrb*$^{\mathrm{Eos/Eos}}$ and *Glrb*$^{\mathrm{Eos/WT}}$ animals using the primers *1*, *2* and *3*. (B) Splicing and transcript expression. The mEos4b sequence is inserted after the signal peptide (shown in gray) before the extracellular domain of the GlyRβ subunit. Right panel: Semi-quantitative RT-PCR. Mixing of spinal cord mRNA from *Glrb*$^{\mathrm{Eos/Eos}}$ and *Glrb*$^{\mathrm{WT/WT}}$ animals at a 1:1 ratio and amplification with primers *4* and *5* produces a PCR pattern that matches the amplification of the two alleles from the heterozygous *Glrb*$^{\mathrm{Eos/WT}}$ mRNA.

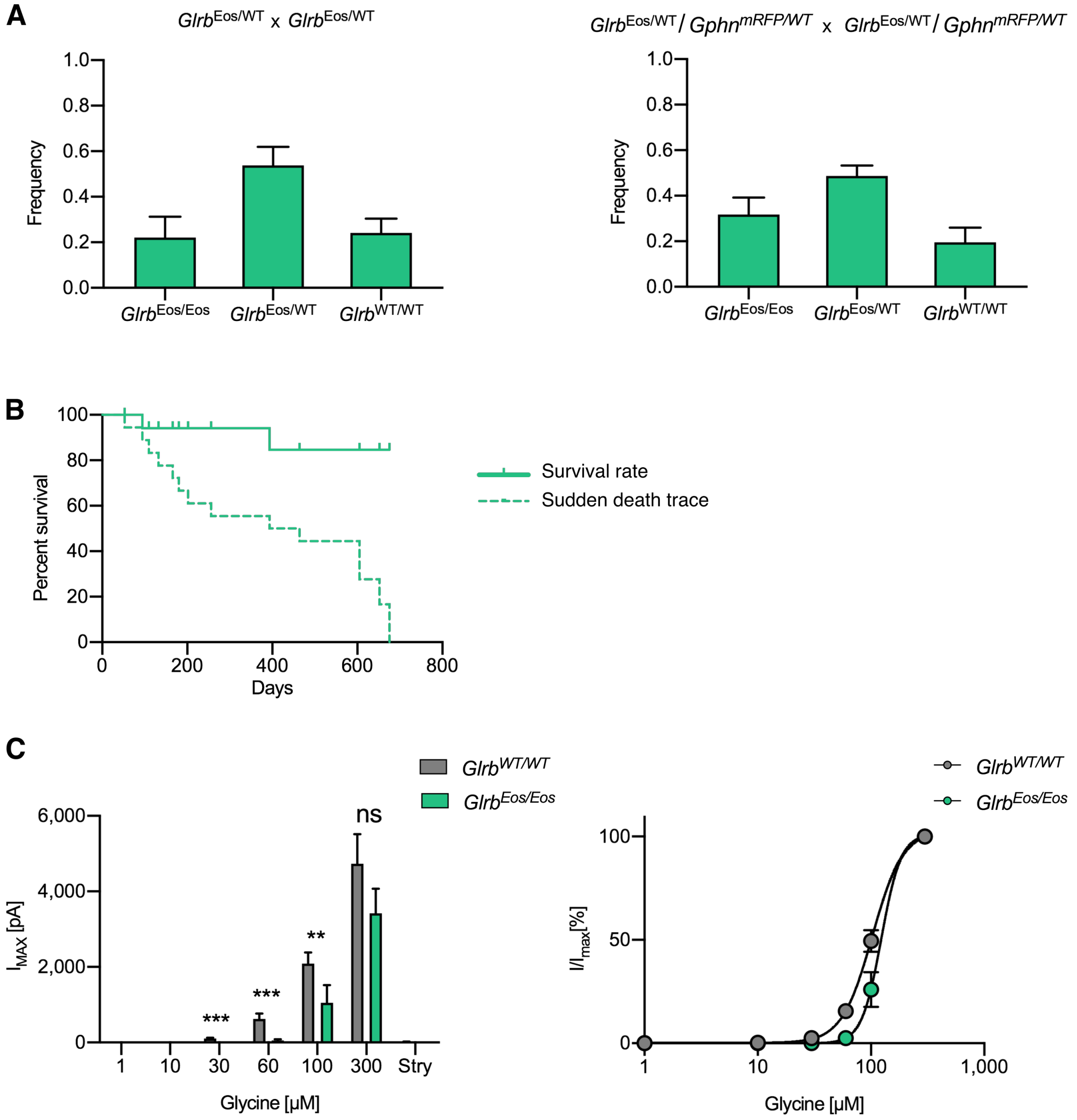


**Fig. S2. Physiological and functional characterization of mEos4b-GlyRβ knock-in mice.** (A) Mendelian inheritance of *Glrb*$^{Eos/Eos}$, *Glrb*$^{Eos/WT}$ and *Glrb*$^{WT/WT}$ genotypes in the offspring of heterozygous single KI matings *Glrb*$^{Eos/WT}$ x *Glrb*$^{Eos/WT}$ (left panel), and heterozygous double KI matings *Glrb*$^{Eos/WT}$/*Gphn*$^{mRFP/WT}$ x *Glrb*$^{Eos/WT}$/*Gphn*$^{mRFP/WT}$ (right panel). Left panel: N =42 pups from 5 litters, plot shows mean ± SEM; right panel: N = 45 pups from 7 litters. (B) Survival plot of homozygous knock-in mEos4b-GlyRβ mice (*Glrb*$^{Eos/Eos}$). N = 19 mice. (C) Left panel: Comparison of mean maximal currents ($I_{max}$) at different concentrations of glycine (1-300 µM) for *Glrb*$^{WT/WT}$ (N = 10) and *Glrb*$^{Eos/Eos}$ (N = 11 neurons). Glycinergic currents were blocked with 10 µM strychnine (Stry) at the end of each recording. Right panel: Normalized dose response curves for *Glrb*$^{WT/WT}$ and *Glrb*$^{Eos/Eos}$ show a subtle shift in the $EC_{50}$ of mEos4b-GlyRβ containing receptors compared to the wild type. Plots show mean ± SEM. **$p < 0.01$, ***$p < 0.001$.

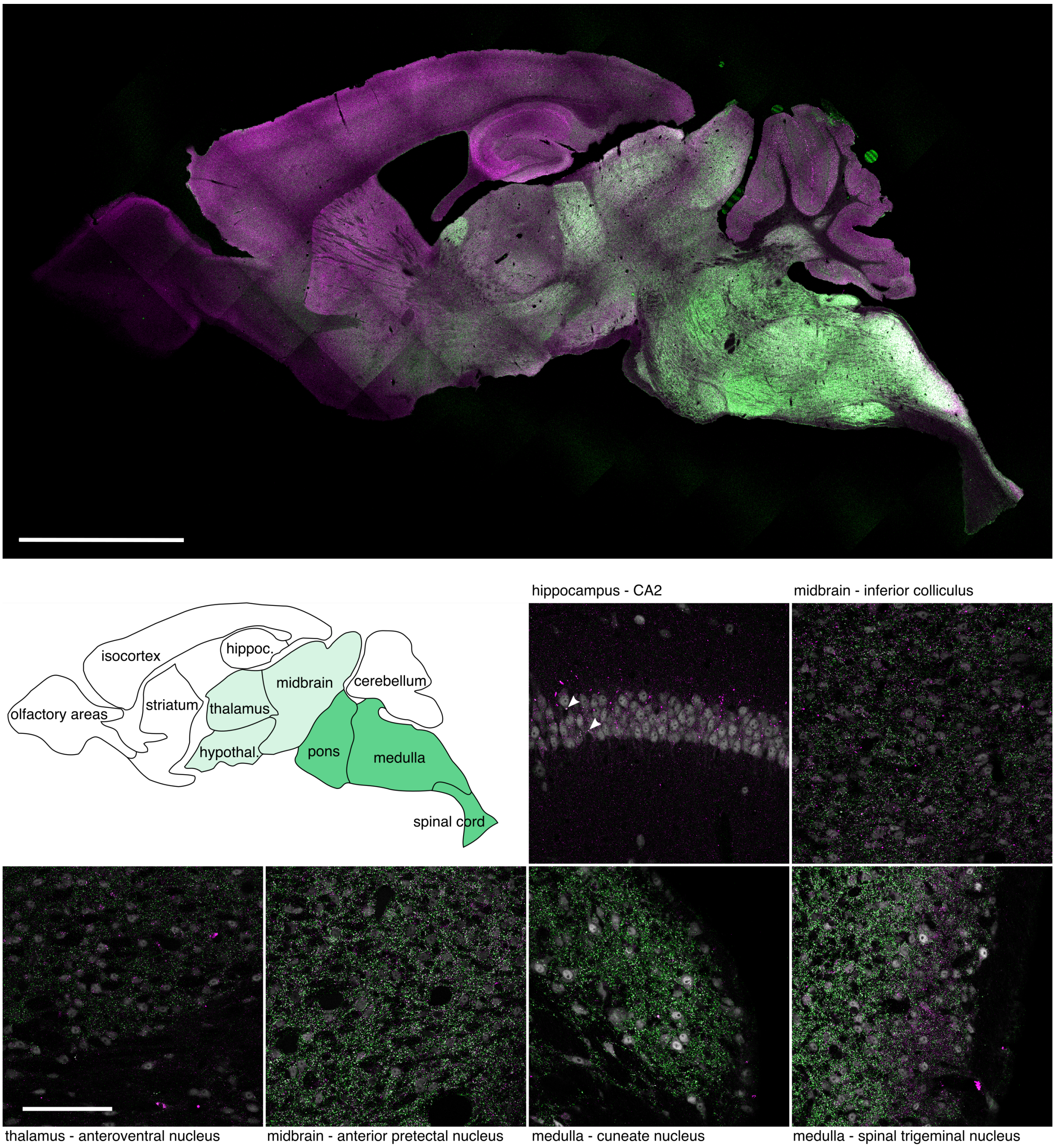


**Fig. S3. Protein expression of mEos4b-GlyRβ in brain sections of knock-in mice.**
Expression of mEos4b-GlyRβ (green) and mRFP-gephyrin (magenta) across brain regions (confocal image of a sagittal vibratome section of a double knock-in *Glrb*$^{Eos/Eos}$ / *Gphn*$^{mRFP/mRFP}$ mouse at 2 months of age). Scale bar = 2.5 mm. The brain atlas depicts the various brain regions with reference to the overall mEos4b-GlyRβ expression levels (white = none-low expression, light green = medium expression, darker green = high expression). High magnification images: Confocal images of various brain regions from the same vibratome section, showing the expression of mEos4b-GlyRβ (green) and mRFP-gephyrin (magenta), as well as NeuN immunolabeling (grey). Arrowheads depict synaptic puncta in the pyramidal cell body layer of the hippocampus that are positive for mEos4b-GlyRβ. Scale bar = 100 μm.

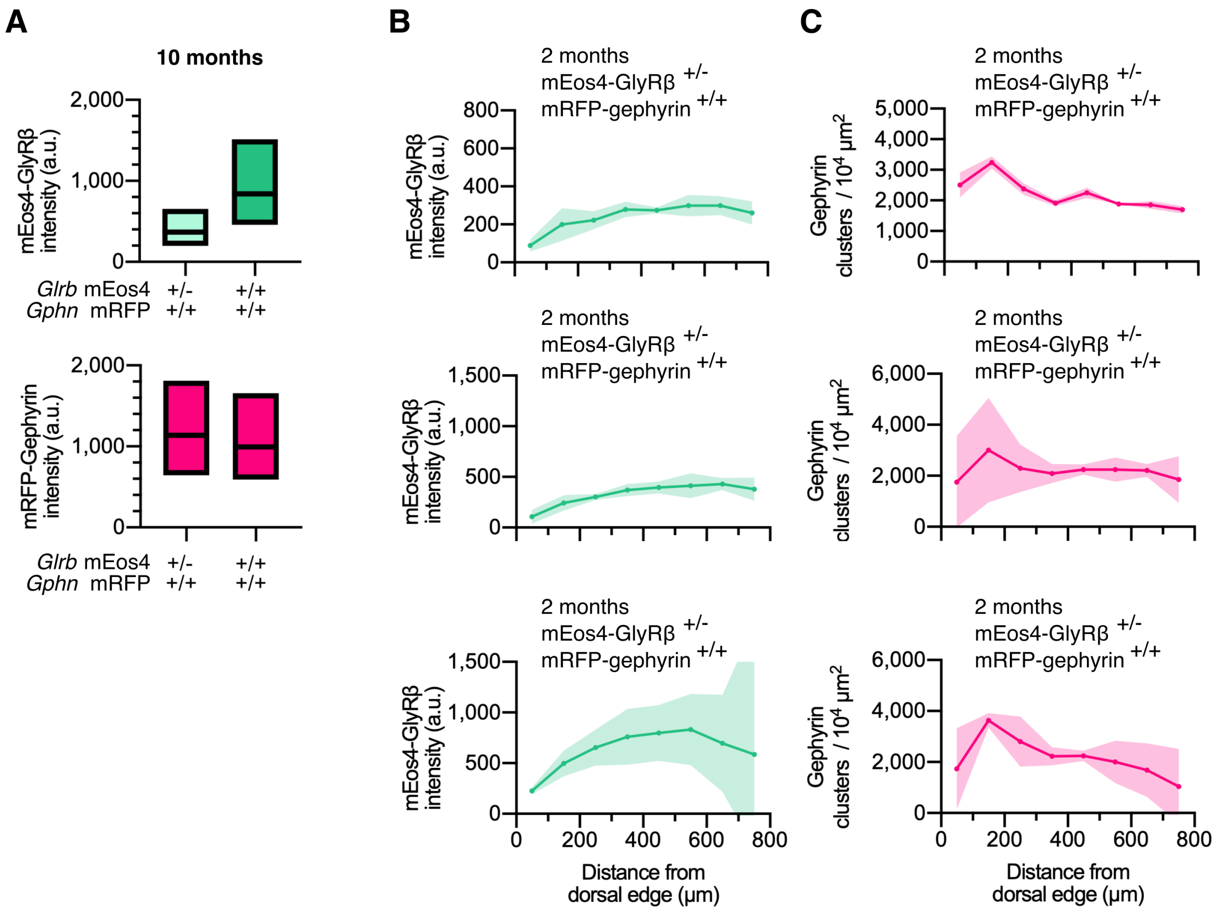


**Fig. S4. Quantitative confocal imaging in 10 month old animals.**

(A) Intensity of mEos4b-GlyRβ and mRFP-gephyrin at spinal cord synapses in homozygous and heterozygous mEos4b-GlyRβ mice, measured in the area indicated by the white square in Fig. 1A. Plots show median and quartiles. N = 5 images per condition from 5 tissue slices per genotype from 2 mice per age group. (B) Quantification of mEos4b-GlyRβ intensity at gephyrin-positive puncta across the spinal cord in 2 month old heterozygous (+/-; *Glrb*$^{\text{Eos/WT}}$) and 10 old month homozygous (+/+; *Glrb*$^{\text{Eos/Eos}}$) and heterozygous animals. Intensities measured in regions as indicated by rectangle in (Fig.1.A). Plots show mean ± 95% confidence interval. N = 2-4 images from 2-4 tissue slices from 2 mice per genotype. (C) Quantification of numbers of gephyrin clusters across the spinal cord in 2 month old heterozygous and 10 month old homozygous and heterozygous animals. Plots show mean ± 95% confidence interval. N = 2-4 images from 2-4 tissue slices from 2 mice per genotype.

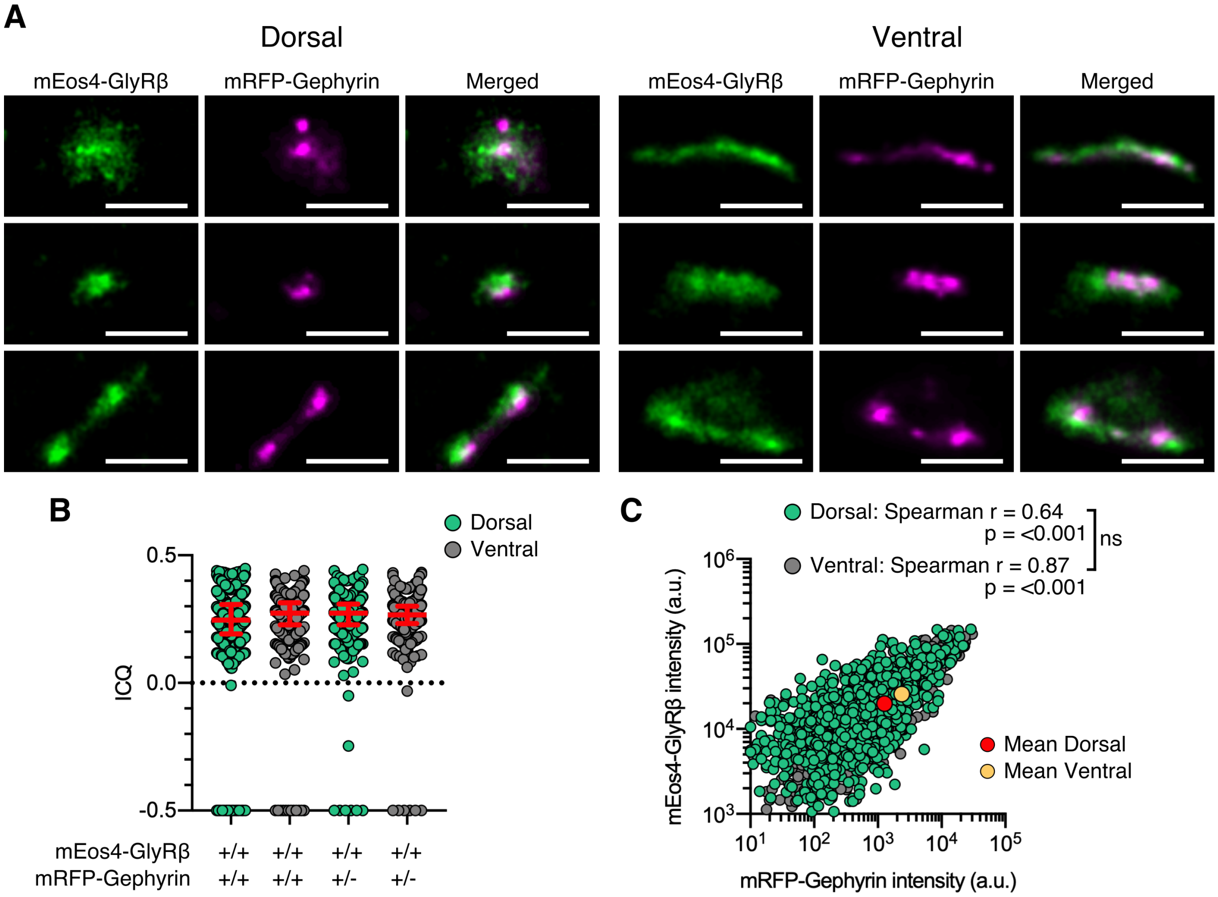


**Fig. S5. Dual-color super-resolution synapse shape & 10 month correlation analysis.**
(A) Examples of synapse shapes and sizes in dorsal and ventral tissue. Scale bar = 500 nm. (B) Intensity correlation quotient (ICQ) of mEos4b-GlyRβ and mRFP-gephyrin in 10 month old heterozygous and homozygous mice. Plot shows median ± interquartile range. N = 466-611 synapses from 17 dorsal and 19 ventral images from 7 tissue slices per spinal cord region from 2 mice per condition. (C) Quantification of GlyR-gephyrin occupancy in 10 month old homozygous mice (*Glrb*$^{Eos/Eos}$ / *Gphn*$^{mRFP/mRFP}$). Non-parametric Spearman's rank shows the same positive correlation at dorsal and ventral synapses. N = 1241 dorsal synapses and 813 ventral synapses. ns = not significant.

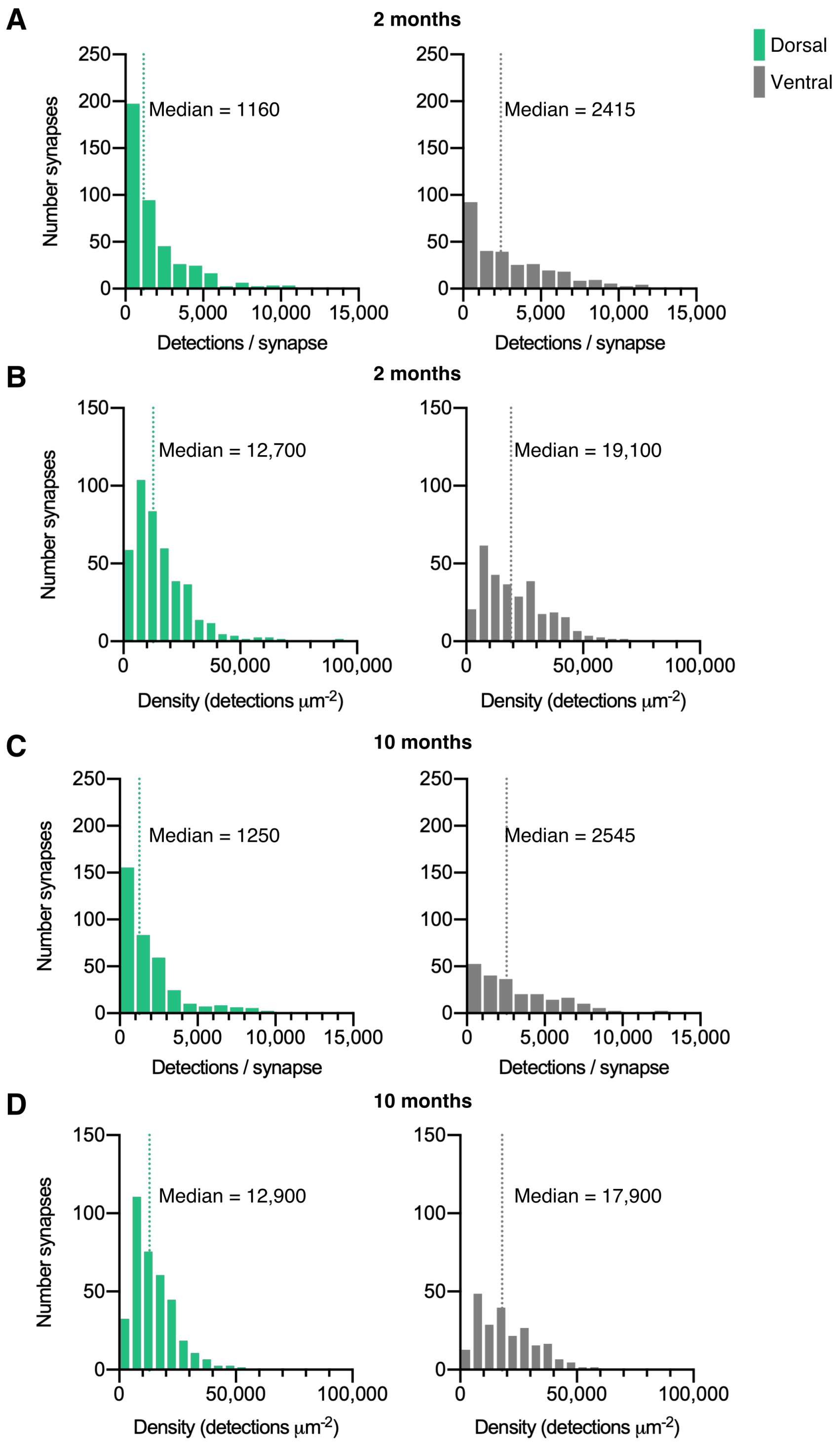


**Fig. S6. Quantification of mEos4b detections at synapses.**
Histograms of mEos4b-GlyRβ detections per synapse (A & C) and density of detections (B & D). N = 433 dorsal synapses and 304 ventral synapses in 2 month old mice. N = 372 dorsal synapses and 234 ventral synapses in 10 month old mice.

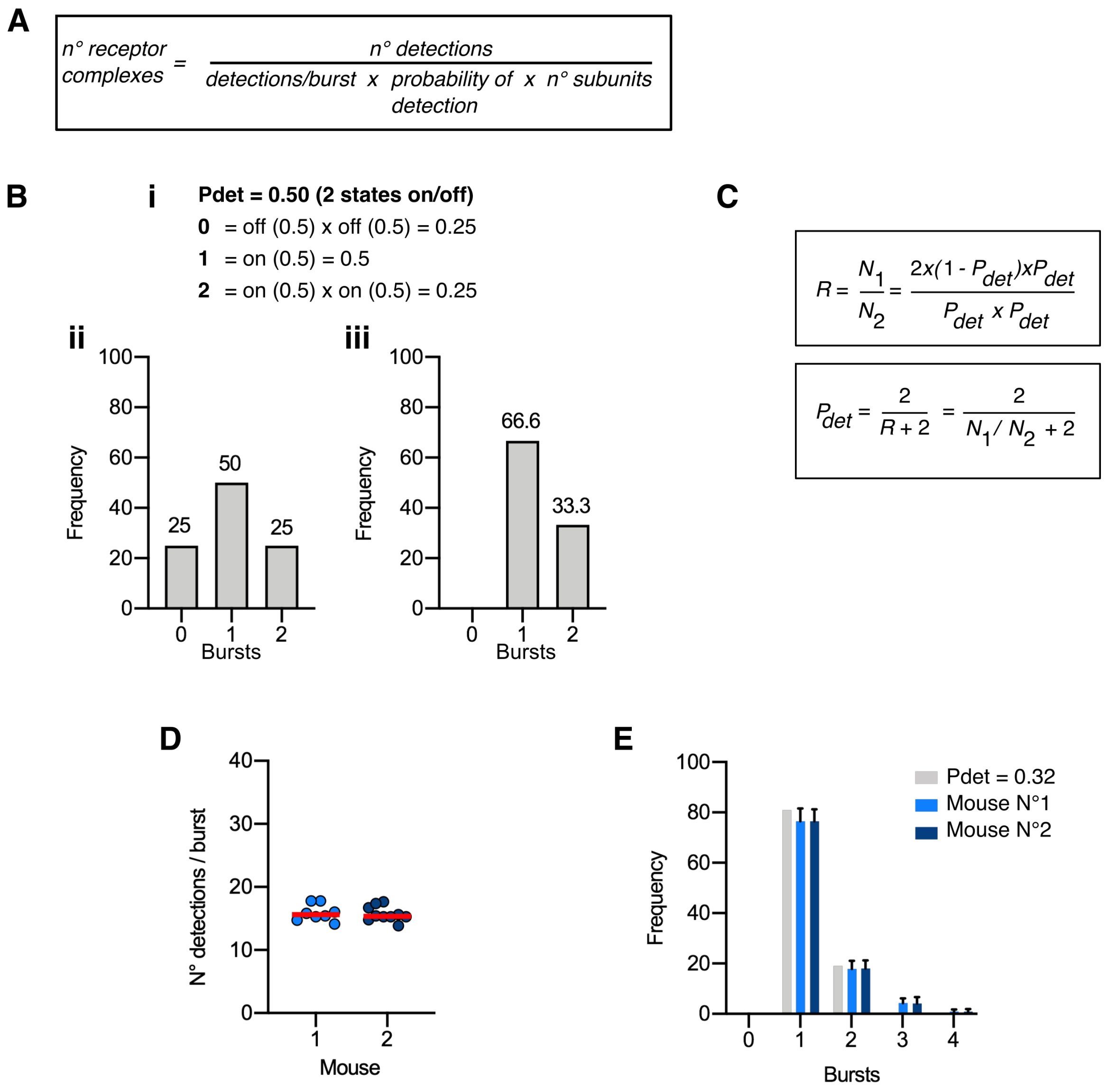


**Fig. S7. Molecule conversion of mEos4b-GlyRβ detections into GlyRs copy numbers.** (A) Descriptive formula for converting fluorophore detections into molecule numbers based on SMLM recordings in sparsely populated, extrasynaptic membrane compartments (Patrizio et al., 2017). Three parameters are required: the number of repetitive detections of the single fluorophores (average detections per burst), the probability that the fluorophore is functional (the probability of detection $P_{det}$), and the stoichiometry of the heteropentameric GlyR complex based on a α3:β2 receptor stoichiometry (Durisic et al., 2014, Patrizio et al., 2017). (B) Theoretical binomial distribution of the number of bursts of fluorescently labeled GlyRβ subunits (imaging 0, 1 or 2 subunits) and a $P_{det} = 0.5$ (i, ii), adjusted for the experimental situation in which only the counts of 1 and 2 bursts per cluster are visible (iii). (C) Derivation of the formula for calculating $P_{det}$, using the counts of 1 and 2 bursts of detections. (D) Number of mEos4b detections per burst in homozygous *Glrb*$^{Eos/Eos}$ animals from our experimental data. N = 8-9 images per mouse from 2 mice. (E) Distribution of the number of bursts of mEos4b detections in homozygous *Glrb*$^{Eos/Eos}$ mice. N = 8-9 images per mouse from 2 mice. The light gray bars represent the adjusted distribution for $P_{det} = 0.32$ that was calculated from the experimental data (blue bars) according to the formula given in (C).

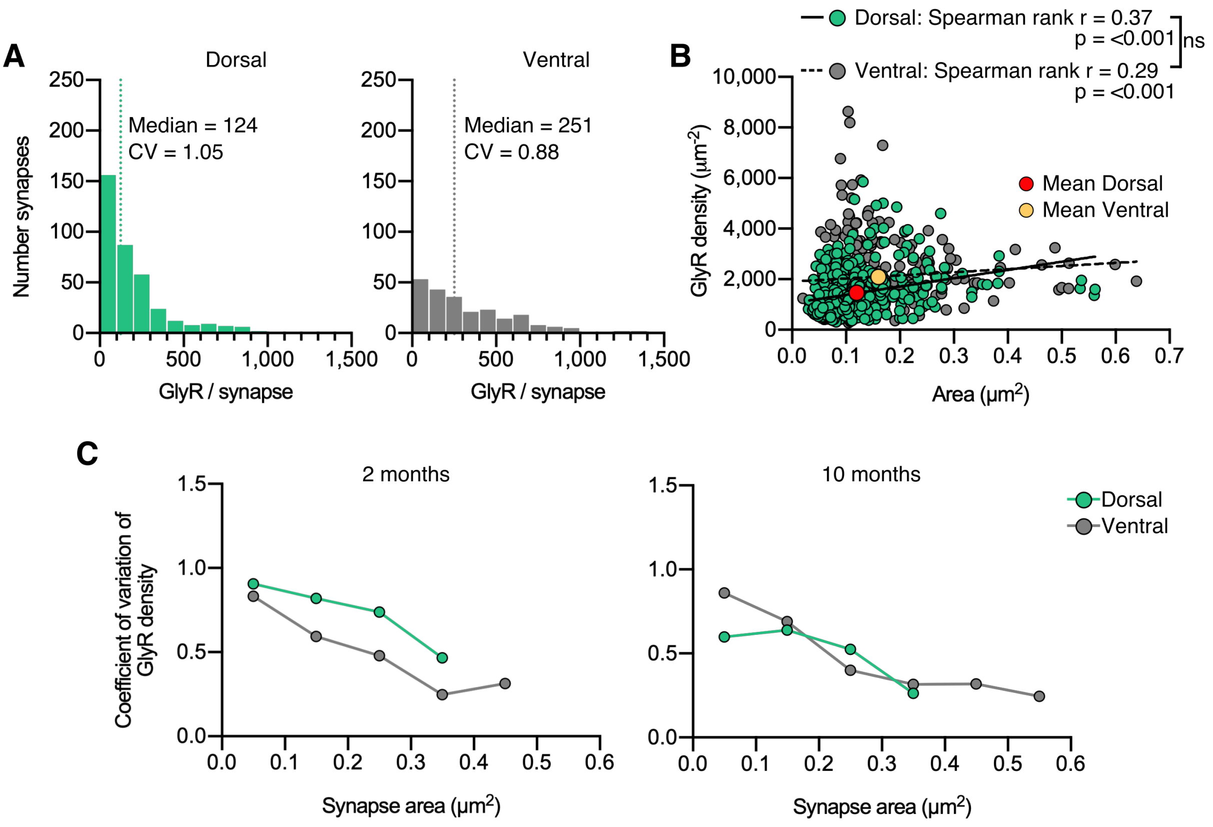


**Fig. S8. Copy numbers and GlyRs densities at synapses (10 months).**
(A) Histogram of the number of GlyRs per synapse calculated from the molecular conversion of detections in 10 month old mice (see Fig. S6 and S7). N = 372 dorsal and 234 ventral synapses from 20 images from 7 tissue slices per region from 2 mice. CV = coefficient of variation. (B) Scatter plot of GlyR density vs synapse area shows no difference between dorsal and ventral synapse densities. ns = not significant. (C) Analysis of the coefficient of variation of GlyR density with respect to synapse area for dorsal and ventral synapses in 2 and 10 month old mice.

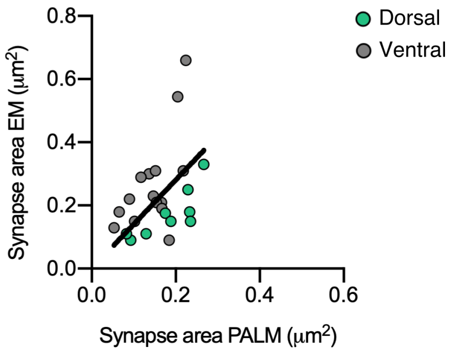


**Fig. S9. Comparison of PALM and EM area measurements.**
Comparison of synapse areas measured by PALM and EM shows a close correspondence.

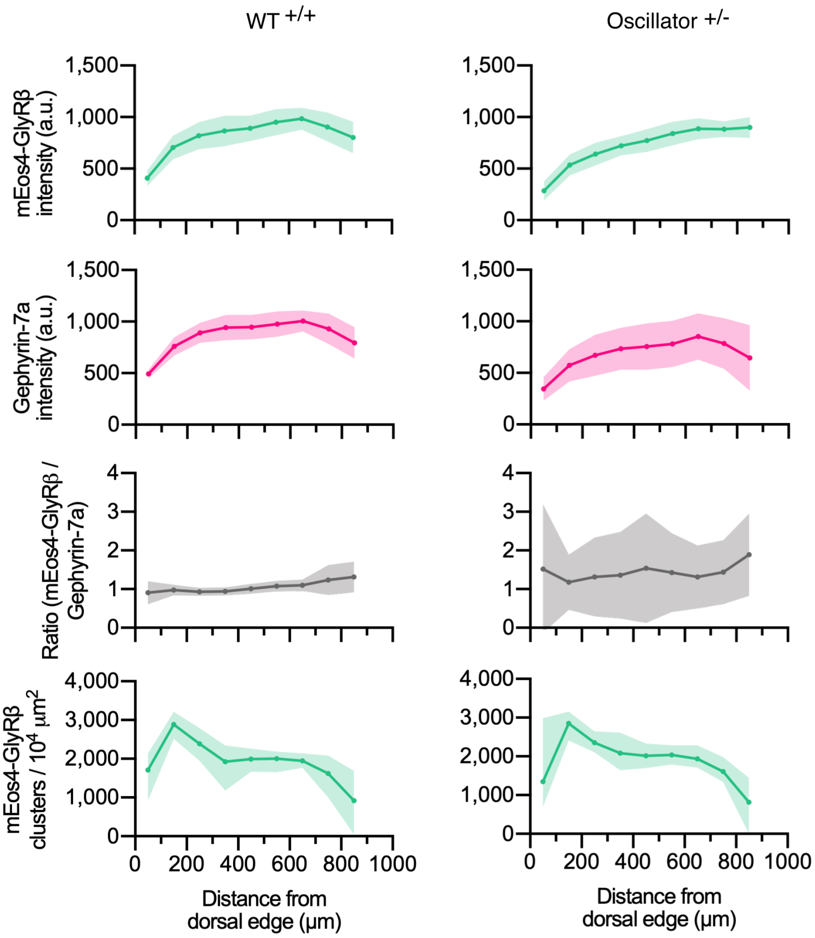


**Fig. S10. Quantitative confocal analysis at mEos4 puncta of the *oscillator* mouse model.** Intensity of mEos4b-GlyRβ and mRFP-gephyrin at mEos4-positive puncta across the spinal cord in mice heterozygous (+/-) for oscillator compared to homozygous WT (+/+) littermates. All mice are homozygous for mEos4b-GlyRβ. Intensities measured in regions as indicated by rectangle in (Fig.1.A). Plots show mean ± 95% confidence interval. N = 9-11 images from 9-11 tissue slices from 2 mice per genotype (same data as in Fig. 4, but using a mask generated in the mEos4b channel instead of the Cy3 immunolabeled gephrin).

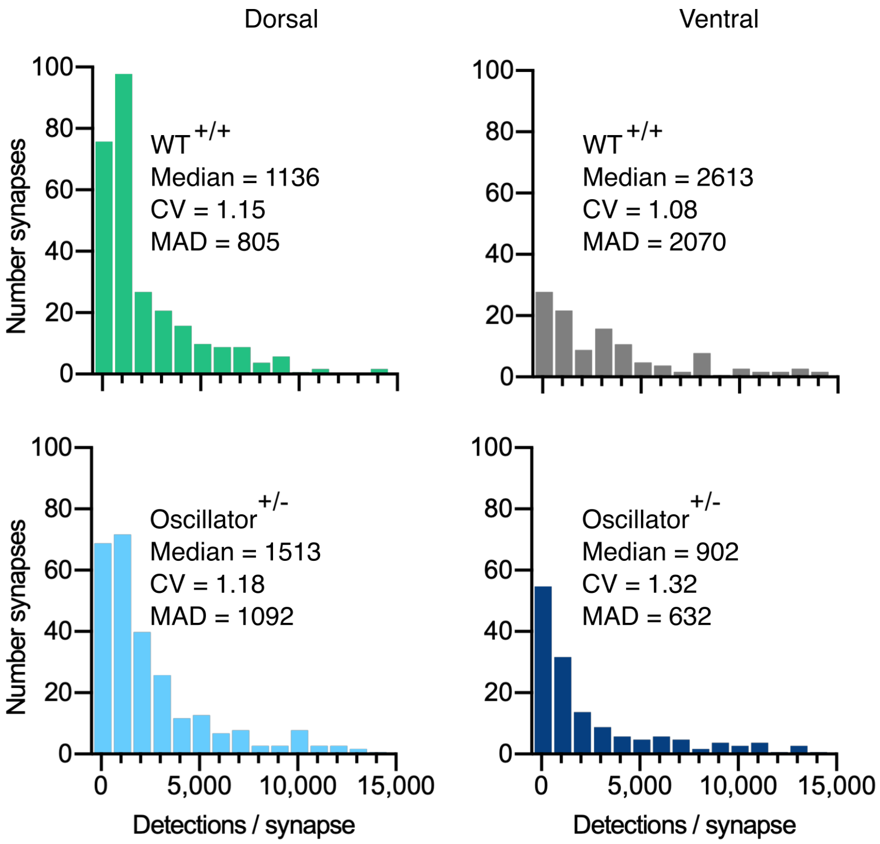


**Fig. S11. Quantification of mEos4b detections at synapses in the *oscillator* mouse model.** Histograms of mEos4b-GlyRβ detections per synapse in heterozygous (+/-) *oscillator* and homozygous (+/+) WT littermates. N = 282 WT dorsal and 120 ventral synapses, 273 *oscillator* dorsal and 156 ventral synapses from 2-4 tissue slices and 2 mice for all conditions. CV = coefficient of variation, MAD = median absolute deviation.

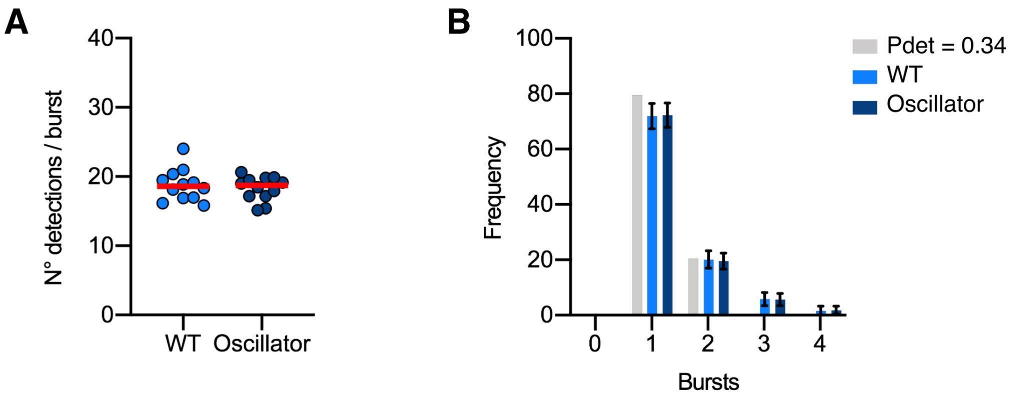


**Fig. S12. Molecule conversion of mEos4b-GlyRβ detections into GlyRs copy numbers in the oscillator mouse model.**
(A) Number of mEos4b detections per burst in homozygous *Glrb*$^{Eos/Eos}$ mice in heterozygous *oscillator* and homozygous WT littermates. N = 12 images per mouse. (B) Frequency distribution of bursts of mEos4b detections in homozygous *Glrb*$^{Eos/Eos}$ mice. The light gray bars indicate a binomial distribution for $P_{det}$ = 0.34 that was calculated from the experimental data as described in Fig. S7. N = 12 images per mouse.
